# Molecular identification of rhizobacterial isolates from *Prosopis limensis* and their effect on the growth of *Raphanus sativus* under salt stress

**DOI:** 10.1101/2021.10.25.464584

**Authors:** Rene Flores Clavo, Esteban A Valladolid Suyón, Karin Reinoza Farroñan, Cristian Asmat Ortega, Gabriel Zuñiga Valdera, Fabiana Fantinatti Garboggini, Sebastian Iglesias-Osores, Carmen Rosa Carreño Farfán

## Abstract

Soil salinity negatively affects the development of agricultural crops. The utilization of plant growth-promoting rhizobacteria is a biotechnological alternative to mitigate this problem. Rhizobacteria were isolated from the roots and rhizosphere of *Prosopis limensis* Bentham “carob” to identify them and determine their potential as plant growth-promoters under salt stress. First, ACC deaminase activity was determined in Dworkin & Foster (DF) minimum medium with 3.0 mM ACC as a source of nitrogen; besides, tolerance to sodium chloride was determined in Nutrient Broth (NB) supplemented with 10% NaCI. Then, bacteria displaying ACC deaminase activity and a 10% NaCI tolerance were selected, identified through genomic analysis targeting the 16s rRNA gene, and inoculated into *Raphanus sativus* L. “radish” seeds to determine the effect on germination percentage under salt stress (80 mM NaCI) and, also on indole production and phosphate solubilization. Isolates were also utilized to evaluate their effect on the growth of radish in saline soils. Genomic analysis showed two bacterial isolates from the genus *Pseudomonas* and one from the genus *Bordetella*: Isolate MW604823 was identified as *Pseudomonas* sp.; isolate MW604824, as *Pseudomonas hunanensis;* and isolate MW604826 as *Bordetella muralis*. Thee isolates demonstrated ACC deaminase activity and tolerance to 10% NaCI. Inoculation of the isolates on radish seeds increased germination percentage compared to the control. The isolates displayed *in vitro* indole production and phosphate solubilization capacity. Moreover, the isolates promoted the growth of radish under salt stress conditions, increasing, leave number, root number, aerial, and root biomass, demonstrating their potential as a biofertilizers.

## 1. Introduction

Plant growth-promoting rhizobacteria (PGPR) are rhizobacteria that can enhance plant growth through a wide variety of mechanisms (Bhattacharyya & Jha, 2012). One of these mechanisms under stress situations is the increased activity enzyme 1-aminocyclopropane-1-carboxylic acid (ACC) deaminase, which decreases ethylene concentrations and the ammonium availability in the rhizosphere of the plant (Gupta & Pandey, 2019). Another mechanism is the production of indole-3-acetic acid (IAA), which acts upon growth and many other physiological responses (Jiang et al., 2020). Phosphate-solubilizing rhizobacteria are another group of microorganisms which contribute to plant growth by increasing soluble phosphorus availability (Tang et al., 2020). Most of these microorganisms are native and help the development of their host plants under stress conditions.

The genus Prosopis is widely distributed on the Peruvian coast. *Prosopis pallida* (carob), is a leguminous tree adapted to arid areas, is an autochthonous species of the dry forests of the northern coasts of Peru. Due to its relevance in the rural economy, since it is used as firewood and its fruits are consumed, most of the previous research on carob has focused on food applications (Padrón & Navarro, 2004). It is also used in construction, honey production, living fences, and to improve saline soils (Harris et al., 2003). *P. pallida* is a particularly important on the northern coast of Peru, due to its wide use and wide distribution in saline areas.

There are studies in which growth-promoting bacteria were isolated from *Prosopis*, one of them found microorganisms such as *Alcaligenes, Bacillus, Curtobacterium, and Microbacterium* in *Prosopis laevigata*; these, when inoculated in crops facilitated root development, significantly improved seed germination and root growth (ROMÁN-PONCE et al., 2017). Nitrogen-fixing bacteria are also found in root nodules on *Prosopis alba* (Salto et al., 2019). *Arthrobacter koreensis*,, identified through 16S rDNA analysis, was also found in *Prosopis strombulifera* which produced abscisic acid (ABA), auxins (IAA), gibberellins (GA1, GA3), and jasmonic acid (JA) under adverse environmental conditions such salt stress (Piccoli et al., 2011).

Salinity in desert soils is not a recent problem, since pre-Columbian times it was present in agricultural activities of the towns that settled on the north coast of Peru (Gamboa et al., 2021). 10% of the total arable land is affected by salinity and between 25% and 30% of irrigated land is affected by salt and rendered unproductive (Zaman et al., 2018). Salinity causes a reduction in the production of growth promoters in plants, which makes them less productive. Plant growth-promoters compensate for a salt-induced hormonal decline in plants and stress mitigation as a defense mechanism (Radhakrishnan & Baek, 2017). Salinity affects crops which leads to economic losses in farmers, and research has been carried out to improve the yield of these crops.

In the last decades, there has been a growing interest in a model plant for studies of physiological stress such as *Raphanus sativus*. It been reported that plant growth-promoting bacteria have been tested in this plant using different concentrations of NaCl. The inoculation of *Lactobacillus* sp. and *Pseudomonas putida* increased radicle lengths compared to the non-inoculated *R. sativus* seeds (Abdallah Hussein & Ho Joo, 2018). The objective of this study was to molecularly identify rhizobacterial isolates from *Prosopis limensis* rhizospheric soil samples and to evaluate their effect on the growth of *R. sativus* under salt stress.

## 2. Material and methods

### 2.1. Bacteria isolation

Bacteria were isolated from root and rhizospheric soil samples of *Prosopis limensis* “carob” trees from a forest located in San Jose, Lambayeque, Peru (06°45’35.65” S, 79°57’41.35” W). In the forest of approximately 700 m^2^, 18 *P. limensis* tress of similar phenotype (height of 5.25-6.10 m) were selected and, from each one of those, three samples of roots and rhizospheric soil were collected. For sampling, a circle was delimited at 0.45 m from the base of the stem, in which three equidistant points were marked. Then, the soil was removed to a depth of approximately 0.60 m, enough to reach the lateral roots, from where samples of 0.1 Kg were taken. Samples were collected and kept under refrigeration during transportation to the Laboratorio de Investigación en Biotecnología Microbiana from the Universidad Nacional Pedro Ruiz Gallo in Lambayeque, Peru and stored at 10±1 °C. Then, phenotypically distinct bacteria were isolated by serial dilutions and cultured in Nutrient Agar plates with 5 % NaCl at 30 °C for up to two days.

### 2.2. PGP traits characterization and salinity tolerance

#### 2.2.1. PGP traits characterization

##### 2.2.1.1. Phosphate solubilization

Phosphate solubilization capacity was determined by the Molybdite technique (Escobar et al., 2011). Bacterial cultures (5 %; 0.25 mL) of each isolate were prepared in 5mL of National Botanical Research Institute’s Phosphate Broth (NBRIP) with 0.85 M of NaCl and incubated at 30 °C daily agitated for 5 min. Each culture was centrifugated at 3000 rpm for 5 min; then, aliquots of the supernatant was placed in tubes. Reaction was considered positive to phosphate solubilization capacity when the solution turned blue, displaying an absorbance of 690 nm. Phosphorous concentrations were determined through spectrophotometry from a standard curve obtained from serial dilutions of 10 ppm of phosphorous. Additionally, phosphate solubilization index (SI) in solid medium was determined (Cecilia Mantilla et al., 2011). To that end, isolates were cultured in Trypticase Soy Agar (TSA) for 24 h, then they were cultured in triplicate in NBRIP agar medium with 1 gL-1 of tricalcium phosphate supplemented with 0.85 M of NaCl. Plates were incubated at 30 °C for 96 h and then, bacterial colony diameter and bacterial inhibition halo were measured to calculate the SI as follows:

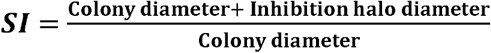

##### 2.2.1.2. IAA production

IAA production. Indole production was evaluated as evidence of in vitro plant growth-promoting traits (Zhou et al., 2017). Bacterial cultures of 9 × 108 cfu/mL were obtained from cultures in 5 mL DF minimum medium with 0.85 M of NaCl at 30 °C for 24 h (Cordova Rojas, 2017). Indole production was determined by the method according to Salkowski (García et al., 2010). Bacterial cultures (5 %; 0.25 mL) of each isolate were prepared in Trypticase Soy Broth (TSB) supplemented with 0.01 gL-1 of tryptophane. After incubation at 30 °C for 72 h, daily agitated for 5 min, each culture was centrifugated at 3000 rpm for 5 min; then, 0.4 mL of each supernatant was placed in tubes and where 1.6 mL of Salkowski reagent was added for 30 min in the dark. Reaction was considered positive to indole production when the solution turned pink, displaying an absorbance of 530 nm. Indole concentrations were determined through spectrophotometry from a standard curve obtained from serial dilutions of 100 ppm of indole acetic acid.

##### 2.2.1.3. ACC deaminase activity

ACC deaminase activity was determined based on a modified qualitative essay utilizing ACC as the only nitrogen source in Dworking & Foster (DF) minimum medium and, also, the same medium but supplemented with ammonium sulfate instead of ACC as a control. After media sterilization in an autoclave, a previously filter-sterilized ACC solution at 0.5 M was added and adjusted to 3.0 mM (Penrose & Glick, 2003; Siddikee, 2010). Isolates were cultured in duplicate in DF minimum medium with ACC and ammonium sulfate for 24 h and, then inoculated in the same media at 30 °C for 24h. Differential turbidity due to the isolates’ growth in the media denoted utilization of ACC as nitrogen source and, thus, positivity for ACC deaminase activity (Angulo et al., 2014).

#### 2.2.2. Salinity tolerance

To evaluate tolerance to salt stress, the isolates were cultured in Nutrient Broth supplemented with 5, 7.5 and 10 % of NaCL at 30 °C. Differential turbidity due to the isolates’ growth in the media denoted tolerance to NaCl (Qin et al., 2014)

### 2.3. Molecular identification of the selected bacterial isolates trough 16S rARN analysis

An isolated colony was used for extraction of genomic DNA, according to the method described by Pospiech et al. (1995), with some modifications (dx.doi.org/10.17504/protocols.io.bpvjmn4n). Amplification of the 16S ribosomal RNA gene was done by means of a polymerase chain reaction (PCR) can be found in (dx.doi.org/10.17504/protocols.io.brrmm546). The obtained products were analyzed and visualized in agarose gel electrophoresis, purified with mini columns (GFX PCR DNA & gel band purification kit, GE Healthcare) is described in this protocol (dx.doi.org/10.17504/protocols.io.brzsm76e) in ABI3500XL Series automatic sequencer (Applied Biosystems), according to the manufacturer’s specifications described in this protocol (dx.doi.org/10.17504/protocols.io.brzpm75n); the work protocols previously described in this section can be found in the web server of protocols.io according to the work (Flores et al., 2021).

Partial sequences of the 16S ribosomal RNA gene obtained from each isolate were assembled into a contig and then compared to the sequences of organisms represented in EZBioCloud 16S Database (https://www.ezbiocloud.net/) using the “Identify” service (Yoon et al., 2017), and species assignment were based on closest hits (Jeon et al., 2014). 16S ribosomal RNA gene sequences retrieved from the database and related to the unknown organism gene were selected for alignment in the Clustal X program (Thompson et al., 1997), and phylogenetic analyzes were performed using the Mega version 7.0 program (Kumar et al., 2016). The evolutionary distance matrix was calculated with the model of Kimura-2 parameters (Kimura, 1980), and the phylogenetic tree constructed from the evolutionary distances calculated by the Neighbor-Joining method (Saitou & Nei, 1987), with bootstrap values from 1000 resampling.

### 2.4. PGP traits of the selected bacteria under NaCl stress

The PGP properties of the selected bacteria were evaluated, as described before, in the presence of 5%, 7.5% and 10% of NaCl.

### 2.5. Effects of bacterial inoculation on R. sativus tolerance to NaCl exposure: pot experiments

For germination essays, seeds of *Raphanus sativus* L. var. Champion were used. First, bacterial inoculants of 1.5 × 10^8^ cfu/mL were obtained from cultures in 5 mL DF minimum medium with 0.85 M of NaCl (∼5 %) and 3 mM of ACC, constantly agitated once a day for 5 min after 6, 12, 18 and 24 h. Then, forty seeds were placed in plates containing a moisten filter paper at the base and put in stove at 30 °C for twelve days (Contreras & Carreño, 2018).

All Petri plates were set with filtered paper containing a solution of distilled water (control) and a solution of 80 mM of NaCl (CE= 6.94 dSm^-1^); while, seeds were previously sterilized with 3 % sodium hypochlorite for 1 min and washed five time with sterilized distilled water (Orhan, 2016). After, seeds were immersed in sterilized distilled water (control group) and in a 1 mL inoculant solution at 30 °C for 10 h in triplicate (Qin et al., 2014). Another control group in which seeds were not inoculated with the rhizobacterial isolates was considered for germination essays. Finally, plates were covered in aluminum foil and kept at 30 °C and germination percentage was determined twelve days after inoculation.

### 2.6. Statistical analysis

All data of germination percentage and rate, leaf number, aerial and root biomass were statistically analyzed in Microsoft Excel^®^. ANOVA test was used to determine statistically significant differences between the means, and Tukey’s Test post-hoc analysis was used to compare the means of all treatments (Hernandez Sampieri et al., 2014).

## 3. Results

### 3.1. Isolation, molecular identification, and phylogenetic analysis of selected isolates

From a total of 78 obtained pure cultures isolated from *P. limensis* rhizospheric soils (Table S1), we selected three isolates: isolate 03, 13 and 31, displaying different colony morphology (Figure 1), for further identification (Table 1) and experimentation.

**Table 1.** Isolation and, molecular identification of rhizobacterial isolates. Closest EZBioCloud (https://www.ezbiocloud.net/) type strains’ accession numbers and their similarity percentages are shown.

| Phenotypic profile |  |  |  |  |  |  |  |  |  |  |  |  | Geographic location | Closest GenBank accession | Similarity (%) | Molecular ID |
| --- | --- | --- | --- | --- | --- | --- | --- | --- | --- | --- | --- | --- | --- | --- | --- | --- |
| Isolate | Gram staining | Catalase test | Oxidase test | TSI Agar Test | Lysine decarboxylation | Citrate | Indol production | Lactose acidity | Manitol acidity | Ornithine acidity | Sorbitol acidity | Voges-Proskauer test |  |  |  |  |
| MW604823 | - | + | + | K/K | - | + | - | - | - | - | - | - | 6°44'13" S<br>79°56'35" W | LBME01000002 | 98.76 | *Pseudomonas* sp. |
| MW604824 | - | + | + | K/K | - | + | - | - | - | - | - | - | 6°44'05"S<br>79°56'58"W | JX545210 | 99.29 | *Pseudomonas* hunanensis. |
| MW604826 | - | + | + | K/K | - | + | - | - | - | - | - | - | 6°44'13"S,<br>79°56'35"W | LC053647 | 97.76 | *Borderella muralis*. |
| + denotes positivity<br>- denotes negative<br>K: alkaline reaction (red color) |  |  |  |  |  |  |  |  |  |  |  |  |  |  |  |  |

**Figure 1.**
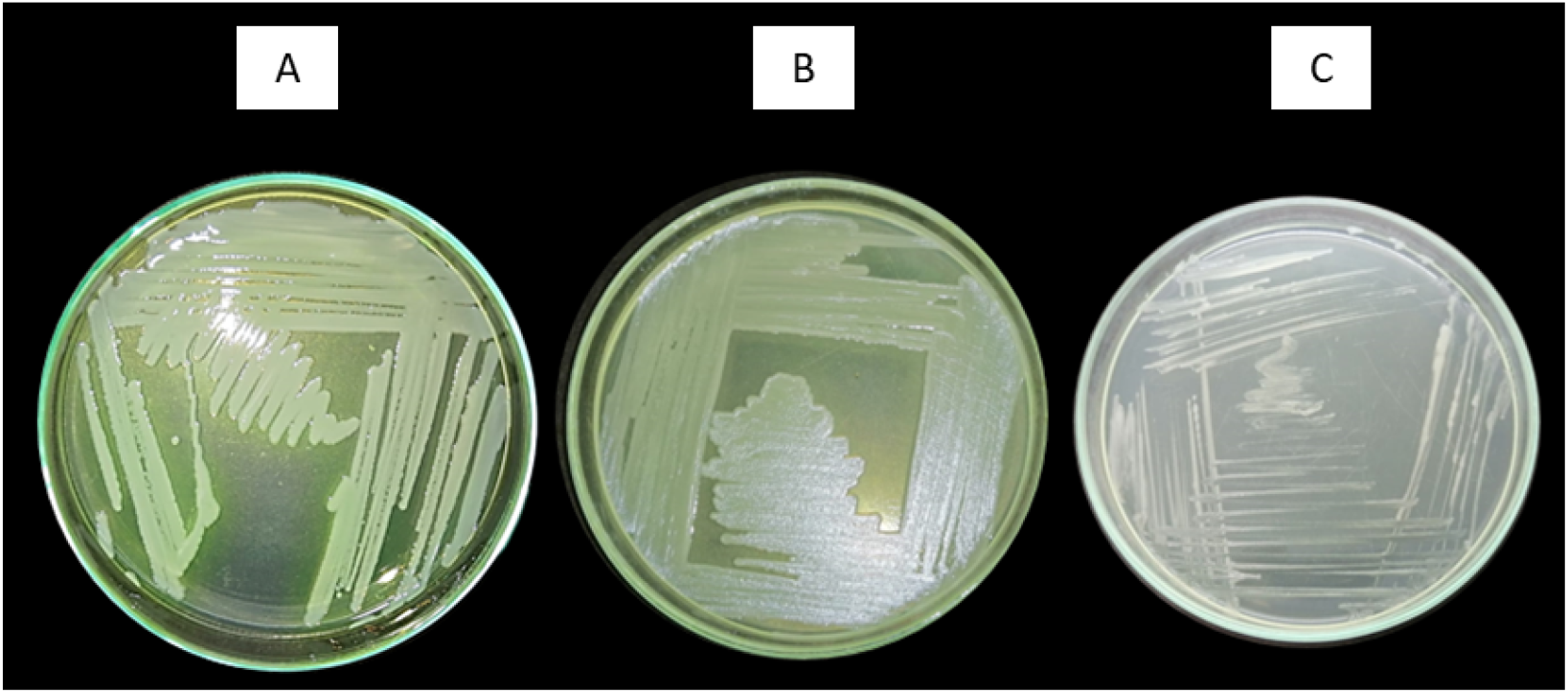
Phenotype of rhizobacterial isolates cultured on agar plates. From left to right A. Isolate 03 (MW604823). B. Isolate 13 (MW604824). C. Isolate 31(MW604826).

Through 16s gene analysis, we identified isolates 03 and 13 as *Pseudomonas* and isolate 31 as *Bordetella*, which were deposited in the GenBank database (GenBank_nih.gov) from the National Center for Biotechnology Information (NCBI) and identified with the accessions numbers: MW604823, MW604824 and MW604826, respectively (Table 1).

Isolate MW604823 was identified as Pseudomonas sp., phylogenetically close to Pseudomonas LBME_s^T^ LBME01000002; isolate MW604824, as Pseudomonas hunanensis, phylogenetically close to Pseudomonas hunanensis LV^T^ JX545210; and isolate MW604826 as Bordetella muralis, phylogenetically close to Bordetella muralis T6220-3-2b^T^ LC053647 and Bordetella tumbae T6713-1-3b^T^ LC053656 (Figure 2).

**Figure 2.**
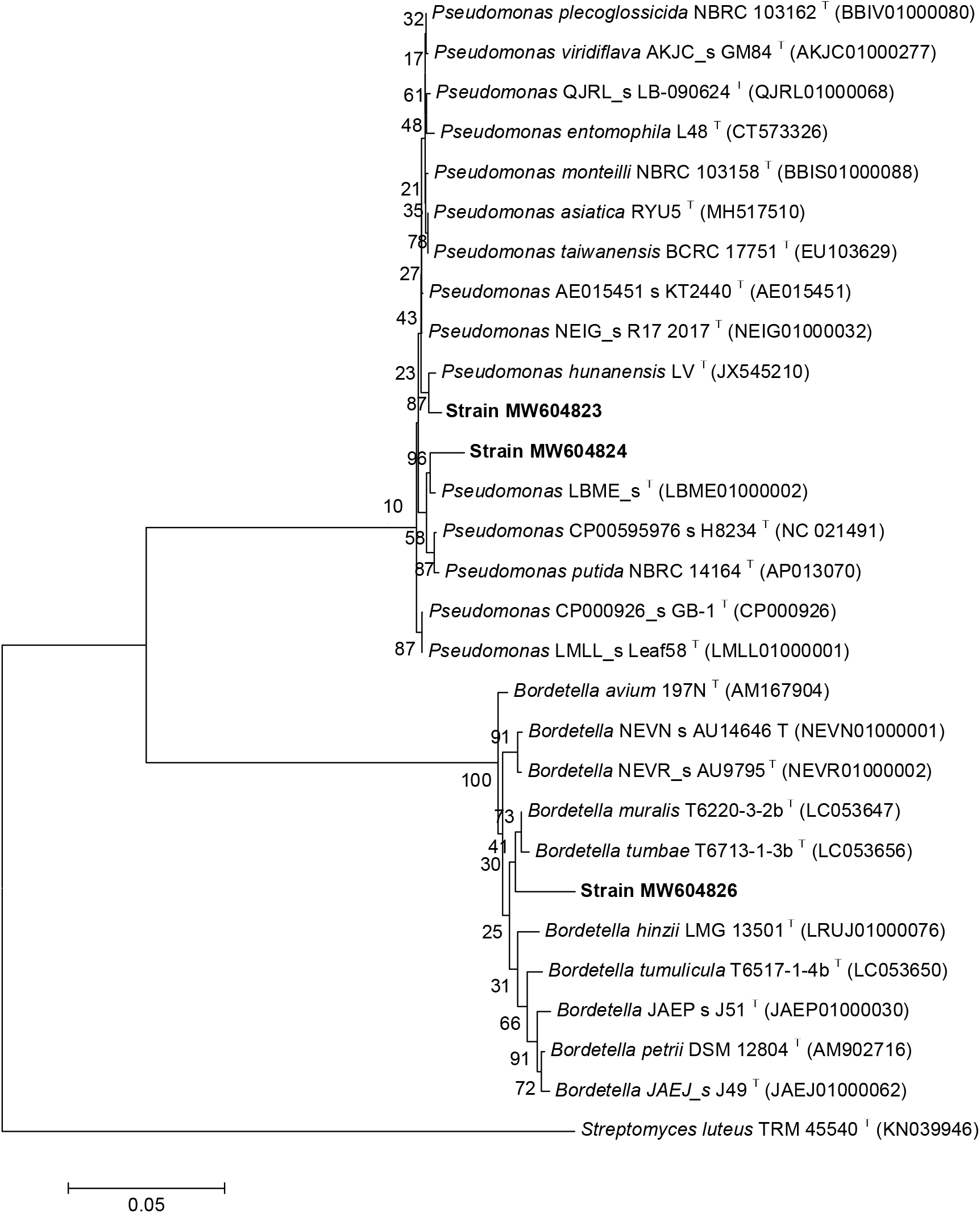
Evolutionary relationships of taxa. The evolutionary history was inferred using the Neighbor-Joining method. The percentage of replicate trees in which the associated taxa clustered together in the bootstrap test (1000 replicates). The tree is drawn to scale 0.05, with branch lengths in the same units as those of the evolutionary distances used to infer the phylogenetic tree. The evolutionary distances were computed using the Kimura 2-parameter method. There were a total of (1200-1500) positions in the final dataset. Evolutionary analyses were conducted in MEGA7.

### 3.2. Tolerance to salt stress, ACC deaminase activity, indole production and phosphate solubilization

All isolates MW604823, MW604824, and MW604826 were able to grow in Nutrient Broth media supplemented with 5, 7.5 and 10 % of NaCl, therefore, the isolates were considered to tolerate salt stress since they showed differential turbidity in comparison with the control (Figure S2). Also, all isolates showed ACC deaminase activity, indole production and phosphate solubilization capacity as shown in Table 2. (Fig. S2, S3 and S4).

**Table 2.**
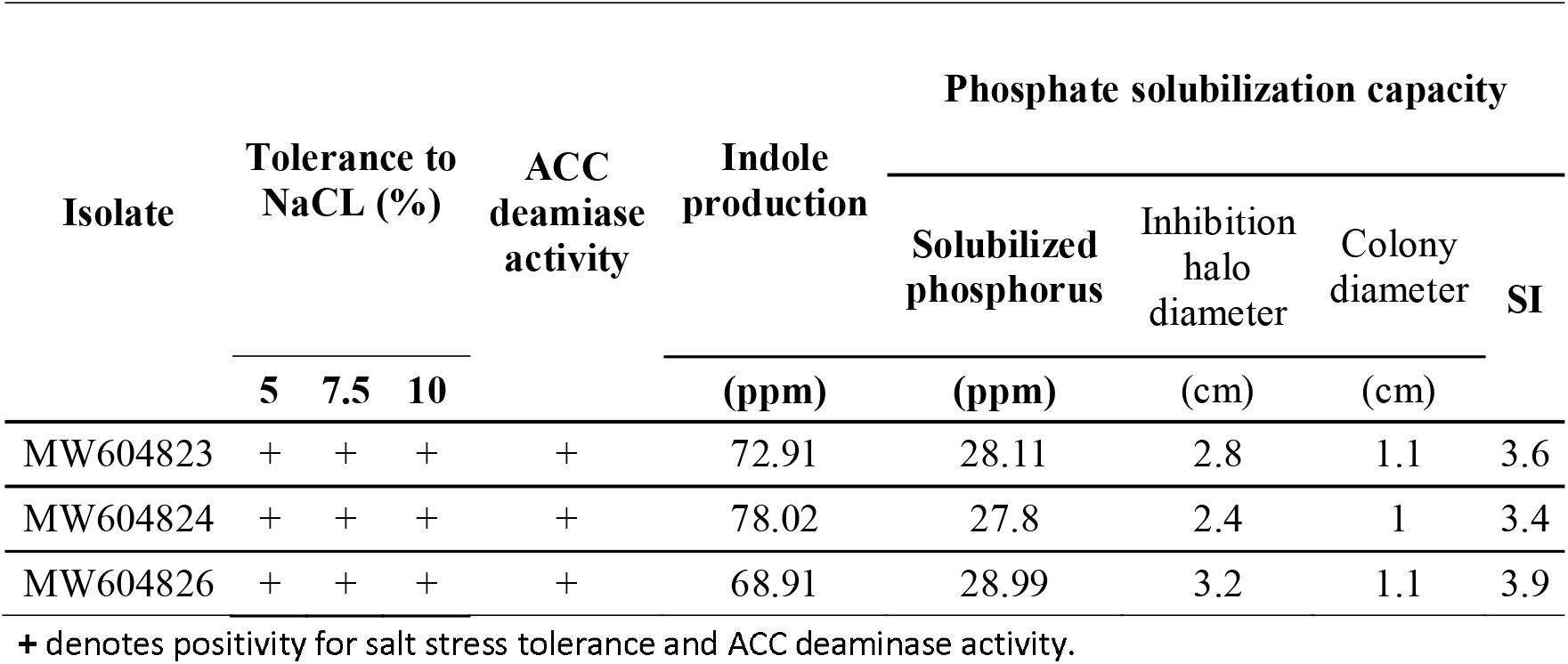
Tolerance to salt stress, ACC deaminase activity, indole production and phosphate solubilization of rhizobacterial isolates.

### 3.3. Isolates’ effect on R. sativus germination

All isolates MW604823, MW604824, and MW604826 significantly increased the germination percentage (Figure 3) from 36 % (control) up to 84.17 %, 82.50 % and 80.00 %, respectively (Table 3).

**Table 3.** Germination percentage of inoculated *R. sativus* seeds.

| Isolate | Germination (%) |
| --- | --- |
| Control | 36.67 (± 5.77) a |
| MW604823 | 84.17 (± 5.20) b |
| MW604824 | 82.50 (± 4.33) b |
| MW604826 | 80.00 (± 4.33) b |

**Figure 3.**
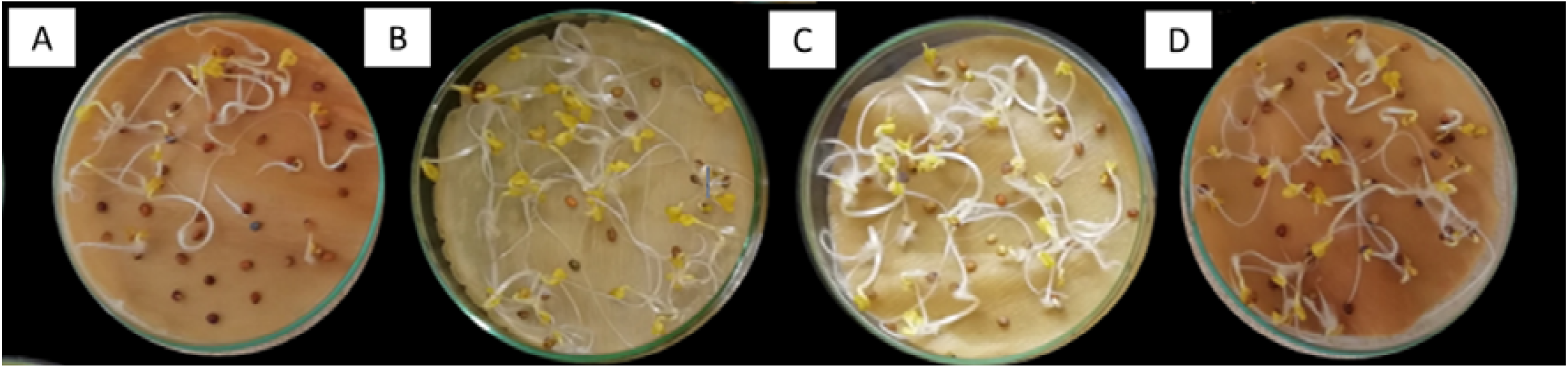
Germinated seeds of inoculated R. sativus. From left to right A. Control. B. Isolate MW604823. C. Isolate MW604824. D. Isolate MW604826.

Means were calculated from three replicates twelve days after inoculation. Means followed by different letters are significantly different according to Tukey’s test with a confidence level of 95 % (Table S2).

### 3.4. Isolates’ effect on the growth of R. sativus under salts stress

All isolates MW604823, MW604824, and MW604826 promoted the growth of *R. sativus* thirty days after inoculation under salt stress conditions in comparison to the control (Figure 4).

**Figure 4.**
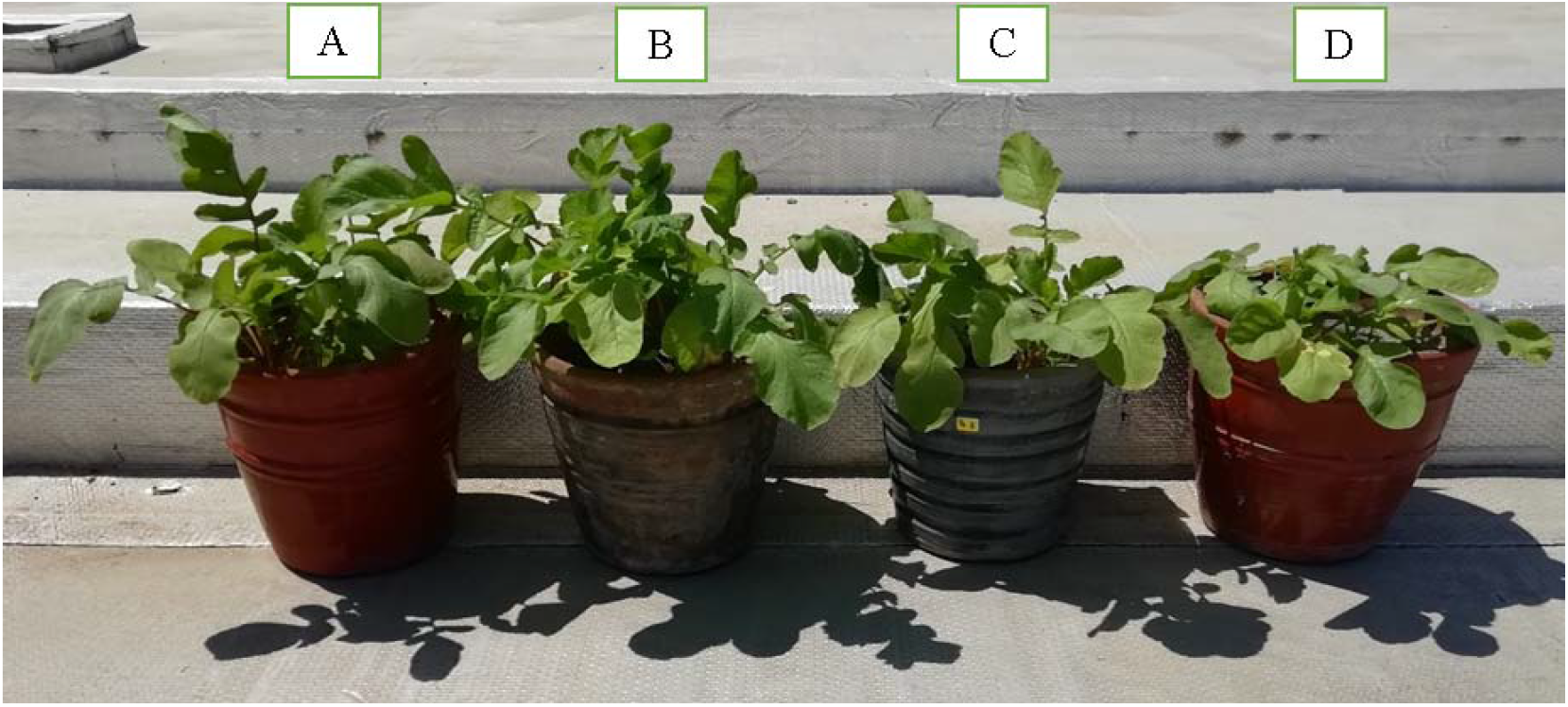
Phenotype of R. sativus plants cultivated in saline soils after thirty days of inoculation. From left to right A. Isolate MW604823. B. Isolate MW604824. C. Isolate MW604826. D. Control.

Among them, isolate MW604824 was the one that most increased the growth of R. sativus considering root biomass (25.40 g). No statistically significant differences were found in plant height of inoculated *R. sativus* plants. Statistically significant differences were found in leaf number, aerial biomass, root number, and root biomass of inoculated *R. sativus* plants (Table 4).

**Table 4.** Plant height, leaf number, aerial biomass, root number and root biomass in inoculated R. sativus seeds.

| Isolate | Plan height (cm) | Leaf number | Aerial biomass (g) | Root number | Root biomass (g) |
| --- | --- | --- | --- | --- | --- |
| Control | 15.69 ($\pm 1.13$) a | 5.42 ($\pm 0.14$) b | 7.56 ($\pm 0.35$) b | 4.83 ($\pm 0.52$) b | 10.47 ($\pm 1.25$) c |
| MW604823 | 20.90 ($\pm 0.90$) a | 6.27 ($\pm 0.29$) ab | 10.18 ($\pm 0.31$) ab | 6.77 ($\pm 0.13$) a | 19.70 ($\pm 1.62$) b |
| MW604824 | 20.99 ($\pm 3.21$) a | 6.37 ($\pm 0.05$) a | 12.51 ($\pm 2.03$) a | 6.78 ($\pm 0.38$) a | 25.40 ($\pm 0.69$) a |
| MW604826 | 19.96 ($\pm 1.06$) a | 5.76 ($\pm 0.59$) ab | 10.29 ($\pm 1.24$) ab | 6.10 ($\pm 0.22$) a | 13.34 ($\pm 1.57$) c |

Means were calculated from three replicates thirty days after sowing. Means followed by different letters are significantly different according to Tukey’s test with a confidence level of 95 % (Table S3).

## 4. Discussion

### 4.1. Characterization of bacterial isolates and PGP traits under salt stress

Salinity stress is an important environmental problem that adversely affects crop production by reducing plant (Hussein & Hoo, 2018). It also microorganisms associated with plants contribute to their growth promotion and salinity tolerance by employing a multitude of macromolecules and pathways, PGPR defence appears to be mediated via increased proline production, enhanced activity of antioxidant enzymes, stimulation of activity of protease and polyphenol oxidases, increased content of phenolics, proteins and chlorophyll (Bano & Muqarab, 2017). In this study, three potential PGPR isolates were selected based on their PGP traits and NaCl tolerance and identified as *Pseudomonas* sp. strain MW604823 of according to its phylogenetic origin, it shares 87% with *Pseudomonas hunanensis* LV^T^ (JX545210), *Pseudomonas* sp. strain MW604824 phylogenetic origin share 96% with *Pseudomonas* LBME_s^T^ (LBME01000002) and *Bordetella* sp. Strain MW604826 too to its phylogenetic origin, it shares 41% with Bordetella tumbae T6713-1-3b^T^ (LC053656) with Bordetella muralis T6220-3-2b^T^ (LC053647), all three grew at different concentrations of salt 5, 7.5 and 10% NaCl (Fig. S1). The isolates obtained from the rhizosphere of *Prosopis limensis* similar to what they reported (Kaushick et al., 2021) this is bacterial communities from the rhizosphere of *Prosopis juliflora* and its native congener *Prosopis cineraria* using high-throughput 16S rRNA gene sequencing, identified predominant phylum proteobacteria in *Prosopis juliflora* while in the rhizosphere of *P. cineraria* it was dominated by the phylum Cyanobacteria, taking into account that *Prosopis limensis* and *Prosopis juliflora* are close because they are native to the American continent and *Prosopis cineraria* their origin is from the Asian continent. Salinity has an overall detrimental effect not only on plants but also on biodiversity and physiology of soil microorganisms (Zhang et al., 2019); the application of PGPR in saline conditions is a promising way to improve agricultural production, the selected bacteria generally maintained the PGP traits (Indol production, phosphate solubilization, ACC deaminase activity) in the presence of saline stress (Table 2).

### 4.2. Effect of bacterial inoculation on Raphanus sativus plant tolerance to NaCl stress

Plant Pots were used for experiments designed to study the effect of the selected bacteria in the tripartite interaction plant-soil-microorganisms, under salt exposure. The studies have focused on the one hand on *Raphanus sativus* (Rabanito) plant growth and oxidative stress and on soil enzyme activities. Inoculation of *R. sativus* plants with bacterial strains resulted increased plant height, leaf number, aerial biomass, root number and root biomass in inoculated *R. sativus* seeds (Table. 4) and (Fig. 4). In addition, the germination percentage in *R. sativus* product of the inoculation with the PGPR resulted in the origin of seeds with a higher growth percentage compared to the control. There are several mechanisms by which PGPRs improve the tolerance of plants to the antioxidant activity of salinity, as the study the (Tirry et al., 2021) of strains identified as *Pseudomonas putida* strain NTM22 and *Pseudomonas cedrina* NTCC12 grow between 2, 4% NaCl and 6% respectively, *Pseudomonas putida* NBRIRA shows in improving the water stress in chickpea, presents increased the plumule length by 1.0 cm at the same level of salinity, however the inoculants used did not increase the pellet growth with 200 mM NaCl, however our results with relevant meanings with respect to control this is microorganisms providing them with the ability to tolerate abiotic stress, given that they present activities such as: ACC deaminase, mineral solubilization, hormone production, biofilm formation, ability to produce siderophores (Kumar et al., 2016). Therefore, it is likely that the enhanced *R. sativus* plants growth and salt tolerance observed in the presence of PGPR microbial inoculants might be primarily attributed to the PGP traits of the bacteria as described above inoculating the plants with the selected bacteria allowed overcoming the negative effects of salinity (Fig. 3).

### 4.3. Effect of bacterial inoculation on soil enzyme activity

It has been reported in similar studies that plant growth-promoting rhizobacteria with ACC deaminase activity isolated from Mediterranean dryland had effects on early nodulation in alfalfa (Cedeño-García et al., 2018), another study found that these rhizobacteria enhanced of the growth and tolerance to salt stress in rice seedlings (*Oryza sativa* L.) through the mediation of ethylene, maintaining ionic homeostasis, improving photosynthetic capacity, and improving the expression of stress-sensitive genes (Ji et al., 2020). The production of indoles mediated by the bacteria isolated in this work can promote plant growth along with the increased synthesis of indole acetic acid (Rupal K et al., 2020). In another study, it has been found that phosphate solubilization can improve cotton growth under alkaline conditions (Ahmad et al., 2018), in comparison to the results that we come to present, it is that the representative microorganisms demonstrating their potential as a biofertilizers for their capacities that they presented in pilot plants of *Raphanus sativus* according to the characteristics in plant height, leaf number aerial biomass, root number, root biomass.

## 5. Conclusions

In this research, the three isolates (MW604823, MW604824, MW604826) showed ACC deaminase activity, indole production, and phosphate solubilization capacity, this is reflected in increased germination rate and percentage, leaf number, aerial biomass, root number, and root biomass in R. sativus.

We recommend carrying out more studies on plant growth-promoting rhizobacteria in native crops of economic interest to the north coast of Peru, to increase the country’s incipient bio-technological capacity. The tolerance to salinity and the phosphorus solubilization characteristics of these bacterial strains which could increase their usefulness as inoculants to improve the productivity of the carob tree in alkaline soil conditions such as those that prevail in the desert areas of Lambayeque, Peru.

These three species reveal to be very evolutionarily distant with the possibility of being new species and in addition to this, their interest in their metabolism is greater because they present important activities for the physiology of the plant and therefore are a source of products for the development of new compounds of biofertilizers.

## Declaration of competing interest

All the authors have no conflict of interest to declare.

## Acknowledgments

This study has been financed by the Concytec - World Bank Project “Improvement and Expansion of the Services of the National System of Science, Technology and Technological Innovation” 8682-PE, to the “World Bank”, to “CONCYTEC” through its executing unit ProCiencia [contract number 190-2018]”

## Appendix A

**Supplementary information**

## Notes

### Competing Interest Statement

The authors have declared no competing interest.

### Summary of Updates

Soil salinity negatively affects the development of agricultural crops. The utilization of plant growth-promoting rhizobacteria is a biotechnological alternative to mitigate this problem. Rhizobacteria were isolated from the roots and rhizosphere of Prosopis limensis Bentham carob to identify them and determine their potential as plant growth-promoters under salt stress. First, ACC deaminase activity was determined in Dworkin & Foster (DF) minimum medium with 3.0 mM ACC as a source of nitrogen; besides, tolerance to sodium chloride was determined in Nutrient Broth (NB) supplemented with 10% NaCI. Then, bacteria displaying ACC deaminase activity and a 10% NaCI tolerance were selected, identified through genomic analysis targeting the 16s rRNA gene, and inoculated into Raphanus sativus L. radish seeds to determine the effect on germination percentage under salt stress (80 mM NaCI) and, also on indole production and phosphate solubilization. Isolates were also utilized to evaluate their effect on the growth of radish in saline soils. Genomic analysis showed two bacterial isolates from the genus Pseudomonas and one from the genus Bordetella: Isolate MW604823 was identified as Pseudomonas sp.; isolate MW604824, as Pseudomonas hunanensis; and isolate MW604826 as Bordetella muralis. Thee isolates demonstrated ACC deaminase activity and tolerance to 10% NaCI. Inoculation of the isolates on radish seeds increased germination percentage compared to the control. The isolates displayed in vitro indole production and phosphate solubilization capacity. Moreover, the isolates promoted the growth of radish under salt stress conditions, increasing, leave number, root number, aerial, and root biomass, demonstrating their potential as a biofertilizers.

