## Supplementary information for "Molecular identification of rhizobacterial isolates from *Prosopis limensis* and their effect on the growth of *Raphanus sativus* under salt stress"

**Appendix A: Supplementary information**

**Table S1.** 78 cultures isolated from *P. limensis* rhizospheric soils.

| Strain  Code | Gram negative bacilli colonies  (N°) | Strains de *Pseudomonas* spp.  N° |
| --- | --- | --- |
| C14 | 6 | 1 |
| C13 | 5 | 3 |
| C10 | 4 | 3 |
| C8 | 3 | 3 |
| C15 | 3 | 2 |
| A17 | 3 | 2 |
| A6 | 2 | 0 |
| B6 | 2 | 0 |
| A7 | 2 | 2 |
| B7 | 2 | 2 |
| C7 | 2 | 0 |
| A9 | 2 | 0 |
| B9 | 2 | 1 |
| C9 | 2 | 2 |
| A10 | 2 | 1 |
| B10 | 2 | 1 |
| B12 | 2 | 2 |
| A13 | 2 | 1 |
| B13 | 2 | 2 |
| B14 | 2 | 1 |
| A15 | 2 | 1 |
| B15 | 2 | 0 |
| A16 | 2 | 2 |
| B16 | 2 | 2 |
| A1 | 1 | 0 |
| B1 | 1 | 0 |
| B4 | 1 | 1 |
| C4 | 1 | 1 |
| B5 | 1 | 0 |
| A8 | 1 | 1 |
| B8 | 1 | 0 |
| A11 | 1 | 0 |
| B11 | 1 | 1 |
| C11 | 1 | 0 |
| A12 | 1 | 0 |
| C12 | 1 | 1 |
| A14 | 1 | 0 |
| C16 | 1 | 0 |
| B17 | 1 | 1 |
| C17 | 1 | 0 |
| B18 | 1 | 1 |
| C18 | 1 | 1 |
| Total 42 | 78 | 42 |

**Table S2**. **Analysis of variance (ANOVA) for germination percentage of inoculated *R. sativus* seeds.** Data were calculated from three replicates twelve days after inoculation ***** shows statistical significance (p ≤ 0.05).

| **Source** | **Sum of squares** | **d.f** | **Mean square** | **F** | **Significance** |
| --- | --- | --- | --- | --- | --- |
| Between groups | 4695.83 | 3 | 1565.28 | 63.94 | * 0.00 |
| Within groups | 195.833 | 8 | 24.4792 |  |  |
| Total | 4891.67 | 11 |  |  |  |

**Table S3**. **Analysis of variance (ANOVA) for Plant height, leaf number, aerial biomass, root number and root biomass in inoculated R. sativus seeds. *** shows statistical significance (p ≤ 0.05).

| **Source** | **Sum of squares** | **d.f** | **Mean square** | **F** | **Significance** |
| --- | --- | --- | --- | --- | --- |
| Plant height | 56.567 | 3 | 18.856 | 5.602 | * 0.023 |
| Leaf number | 1.784 | 3 | 0.595 | 5.185 | * 0.028 |
| Aerial biomass | 36.842 | 3 | 12.281 | 8.343 | * 0.008 |
| Root number | 7.542 | 3 | 2.514 | 20.89 | * 0.000 |
| Root biomass | 400.924 | 3 | 133.641 | 74.477 | * 0.000 |

**Figure S1.** **Isolates’ tolerance to salt stress** All isolates 03, 13, and 31 were able to grow in Nutrient Broth media supplemented with 10 % of NaCl and showed differential turbidity in comparison with the control.

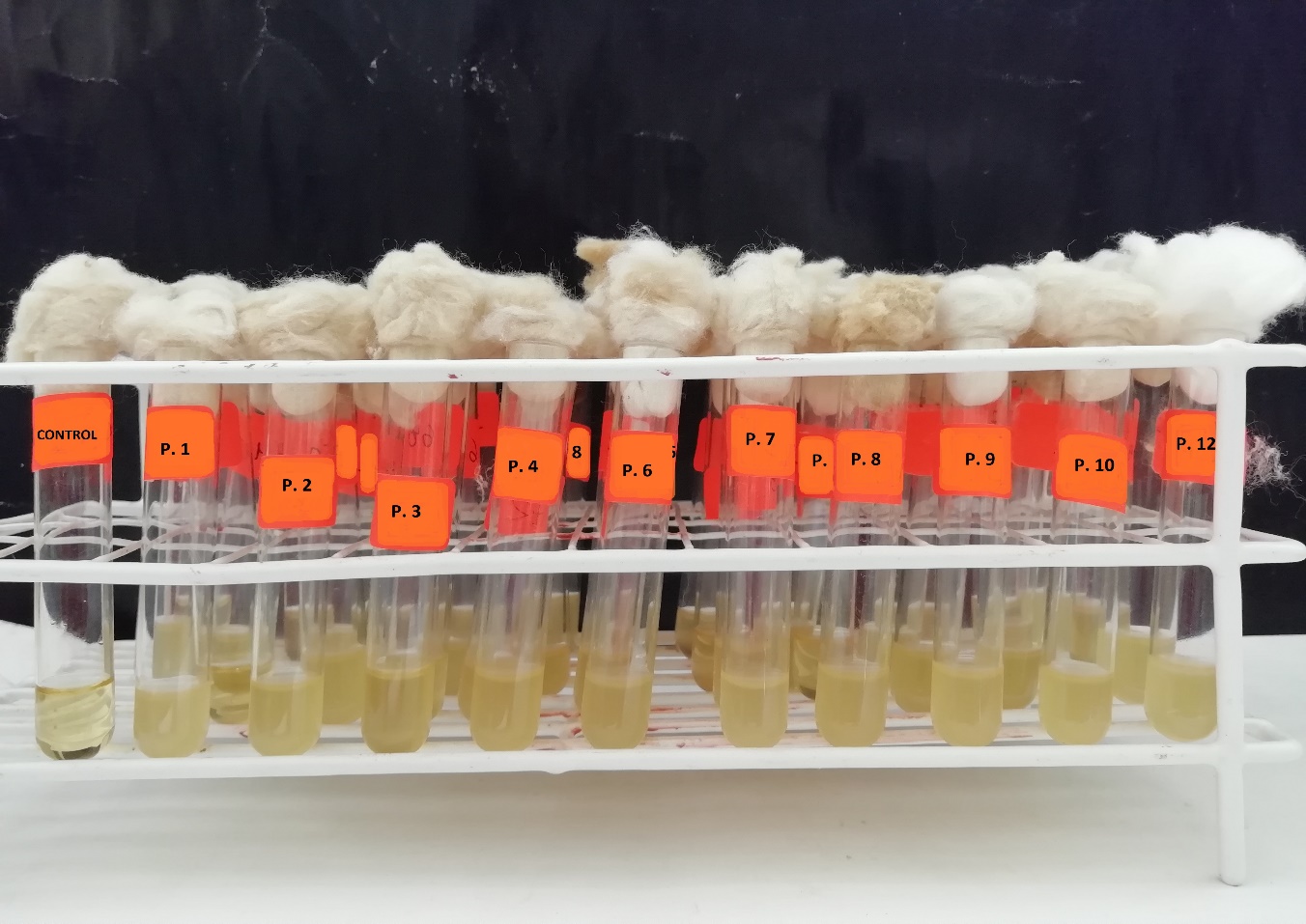

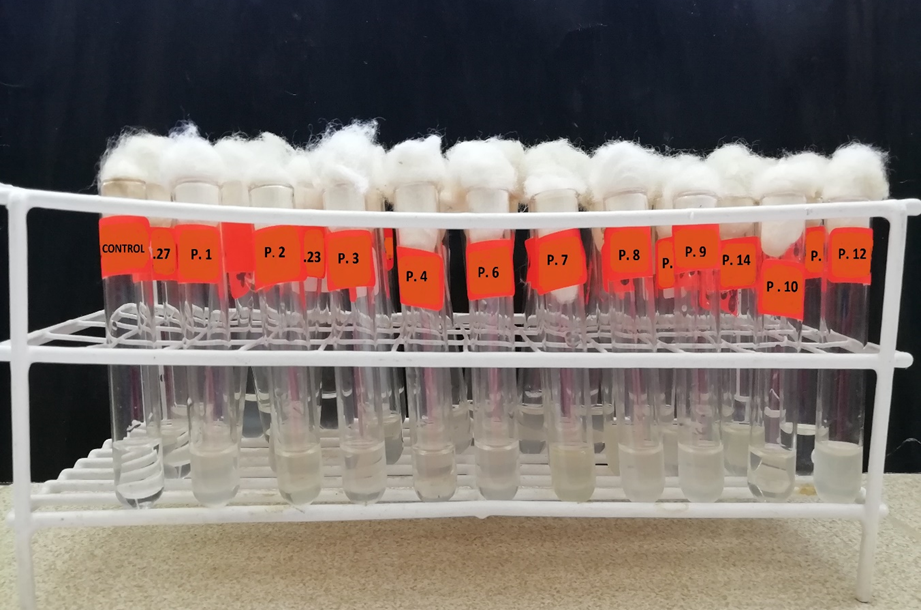
**Figure S2.** **ACC deaminase activity of rhizobacterial isolates.** All isolates 03, 13, and 31 were able to grow in DF media supplemented with 0.5 M of ACC and showed differential turbidity in comparison with the control.

**Figure S3.** **Indole production of rhizobacterial isolates.** All isolates 03, 13, and 31 were able to grow in TSB media supplemented with 0.01 gL^-1^ of tryptophane and showed differential turbidity in comparison with the control.

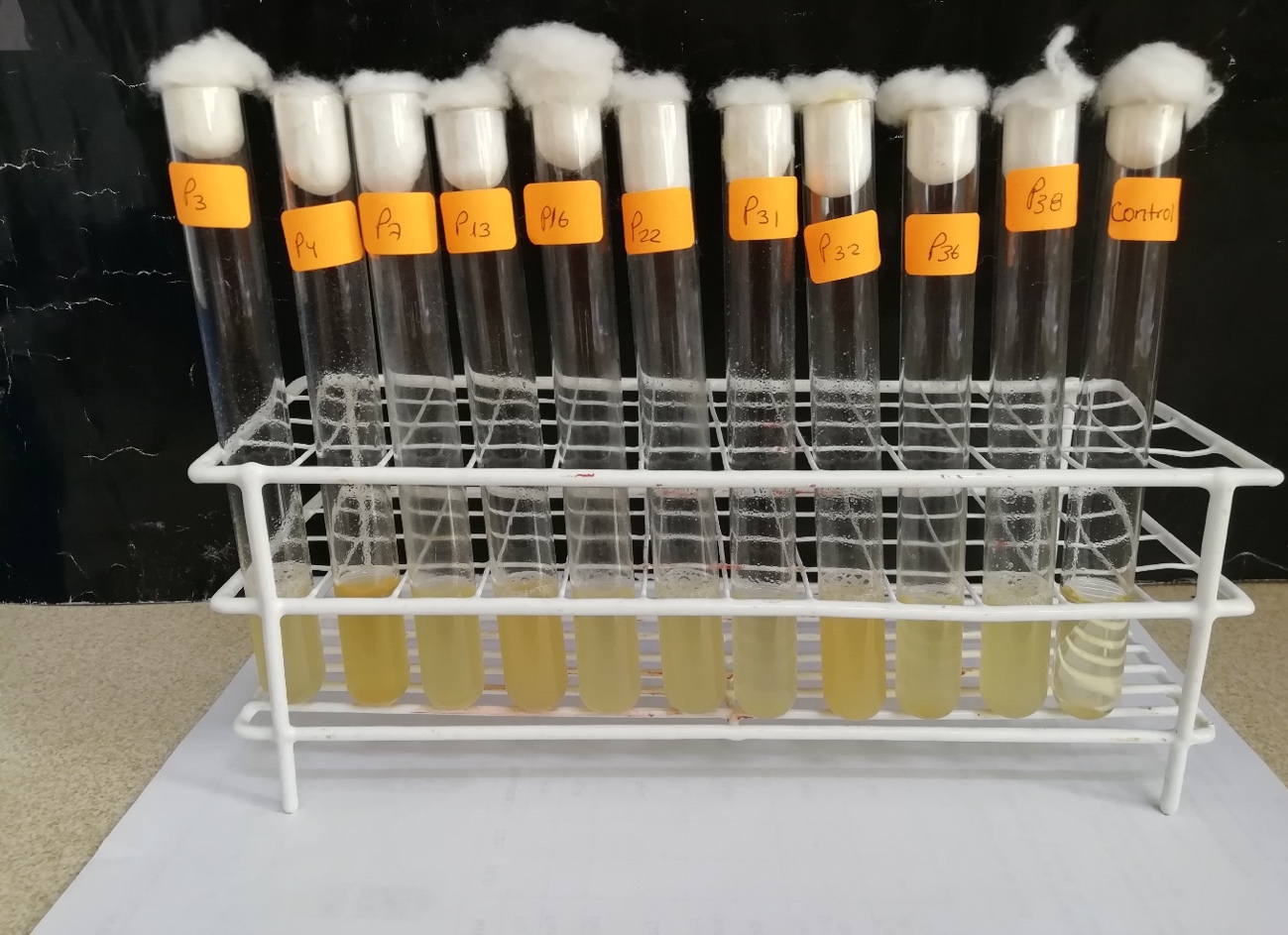

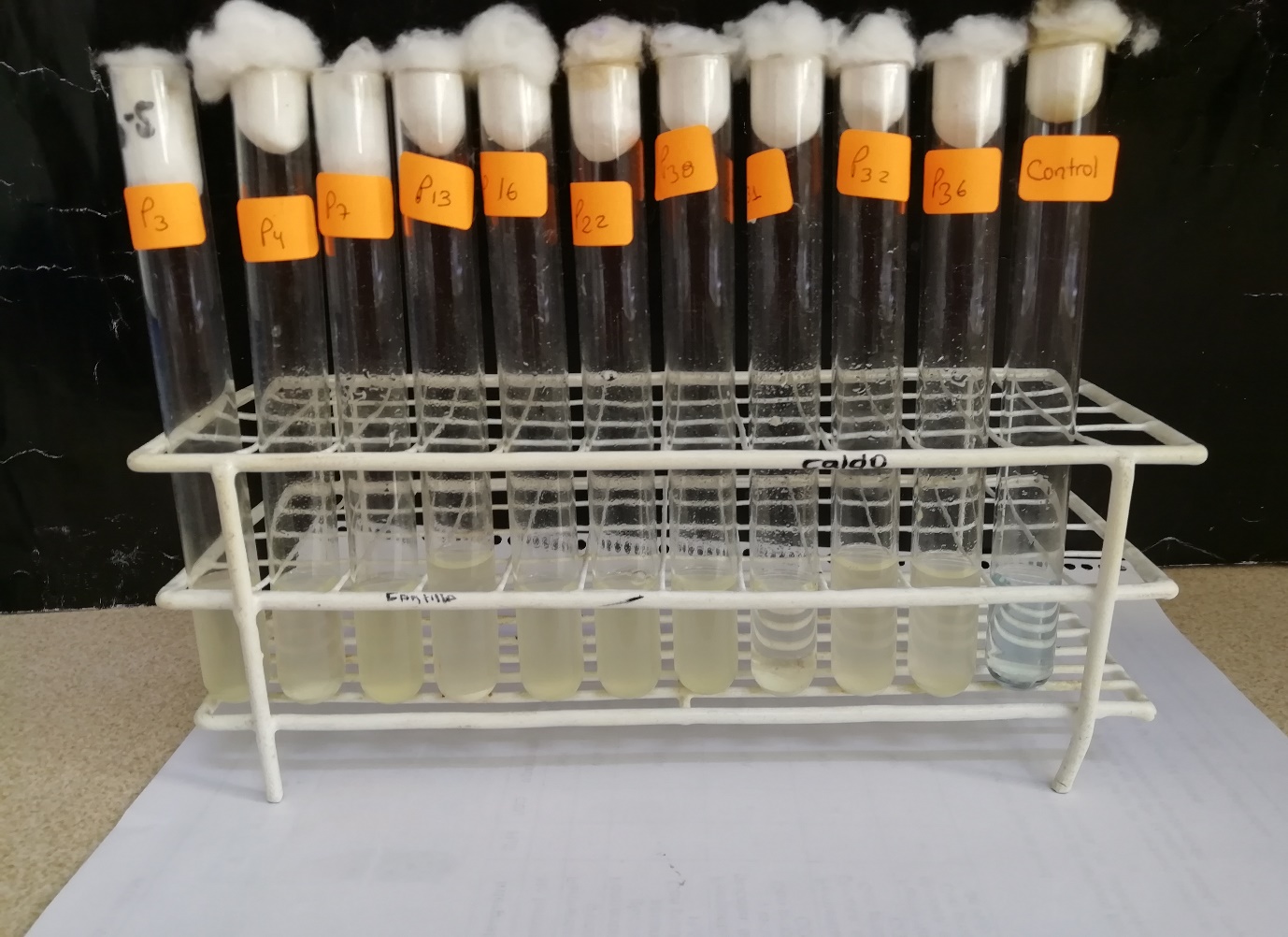

**Figure S4. Phosphate solubilization capacity** **of rhizobacterial isolates.** All isolates 03, 13, and 31 were able to grow in TSB media supplemented with 1 $\mathrm{gL}^{-1}$ of tricalcium phosphate and showed differential turbidity in comparison with the control.
